# Ilastik: a machine learning image analysis platform to interrogate stem cell fate decisions across multiple vertebrate species

**DOI:** 10.1101/2024.12.21.629913

**Authors:** Alma Zuniga Munoz, Kartik Soni, Angela Li, Vedant Lakkundi, Arundati Iyer, Ari Adler, Kathryn Kirkendall, Frank Petrigliano, Bérénice A. Benayoun, Thomas P. Lozito, Albert E. Almada

**Author notes:** Equal Contributors (Alma Zuniga Munoz, Kartik Soni).

## Abstract

Stem cells are the key cellular source for regenerating tissues and organs in vertebrate species. Historically, the investigation of stem cell fate decisions *in vivo* has been assessed in tissue sections using immunohistochemistry (IHC), where a trained user quantifies fluorescent signal in multiple randomly selected images using manual counting—which is prone to inaccuracies, bias, and is very labor intensive. Here, we highlight the performance of a recently developed machine-learning (ML)-based image analysis program called Ilastik using skeletal muscle as a model system. Interestingly, we demonstrate that Ilastik accurately quantifies Paired Box Protein 7 (PAX7)-positive muscle stem cells (MuSCs) before and during the regenerative process in whole muscle sections from mice, humans, axolotl salamanders, and short-lived African turquoise killifish, to a precision that exceeds human capabilities and in a fraction of the time. Overall, Ilastik is a free user-friendly ML-based program that will expedite the analysis of stained tissue sections in vertebrate animals.

## Motivation

Assessing *in vivo* cell fates in vertebrates is commonly done using immunohistochemical analysis on tissue sections. Quantification of desired fluorescent signal is accomplished by measuring fluorescence in several randomly selected images from tissue sections. Ilastik is an ML-based program that allows for the precise quantification of the desired antigen in the entire tissue section, essentially removing user bias, loss of data due to random sampling, and vastly speeding up the analysis process.

## Introduction

Stem cells are the building blocks for regenerating tissues and organs in vertebrate animals. While many factors contribute to the lengthy timelines of stem cell studies in animal models^1^, one of the major impediments to progress is the time it takes to quantify stem cell fates *in vivo* using immunohistochemistry (IHC). Performing IHC on thinly sliced tissue sections is the gold-standard method for assessing cell fate decisions *in vivo*^2,3^. To break down the process, an investigator will harvest desired tissues, section the tissues to a specific thickness, and then use IHC to stain the sections with fluorescently labeled antibodies that detect key markers of cell identity or cell states such as proliferation, differentiation, or cell death. Subsequently, the investigator randomly selects multiple images in a blind fashion followed by quantification of antigen-specific signal in a secondary program like ImageJ/Fiji ^4,5^. However, this analysis workflow creates several problems such as: (1) making broad conclusions from a relatively small number of images being quantified in tissue sections; (2) investigator error when fields of view are randomly selected and later quantified manually for analysis; and not to mention the (3) labor-intensive nature of doing manual counting in programs like ImageJ. Consequently, we suspect that a combination of these issues slows down progress in the field of regenerative biology and likely contributes to the data reproducibility crisis currently ongoing in academia^6–9^.

In recent years, there has been great progress in applying machine learning (ML)-based approaches, a type of artificial intelligence (AI), to expedite the analysis of data-rich images in biomedical research. For example, several ML-based programs have been developed including Imaris (RRID:SCR_007370), Cell Profiler Analyst^10^, Microscopic Image Analyzer (MIA)^11^, CellPose 1 and 2^12,13^, and Ilastik^14^. Conveniently, many of these programs were developed with user-friendly interfaces that eliminate the need for advanced programming skills. Despite the obvious benefits, many labs in regenerative biology continue to perform manual counting of fluorescently marked cells in tissue sections in a small number of images. We suspect that with further education and extensive validation highlighting their potential, investigators will feel confident to transition away from manual counting and embrace a new AI-based future. Therefore, in this study, using the ML-based imaging analysis program, Ilastik, and skeletal muscle as a model system, we demonstrate that Ilastik can precisely quantify PAX7-positive (PAX7+) muscle stem cells (MuSCs) in whole resting and injured skeletal muscle sections in mice, humans, axolotl salamanders, and killifish. Importantly, what would previously take researchers days and weeks to compute manually, Ilastik does it in hours. Collectively, we hope this validation study will speed up the transition from performing manual counting of imaging data to ML-based image analysis workflows.

## Results

### An overview of the Ilastik workflow

Here we present our pipeline for the quantitative analysis of MuSCs in skeletal muscle sections beginning with (A) sample acquisition and (B) pre-processing of images after IHC, followed by (C) Pixel and (D) Object Classification in the Ilastik program (**Figure 1**)^14^.

**Figure 1.**
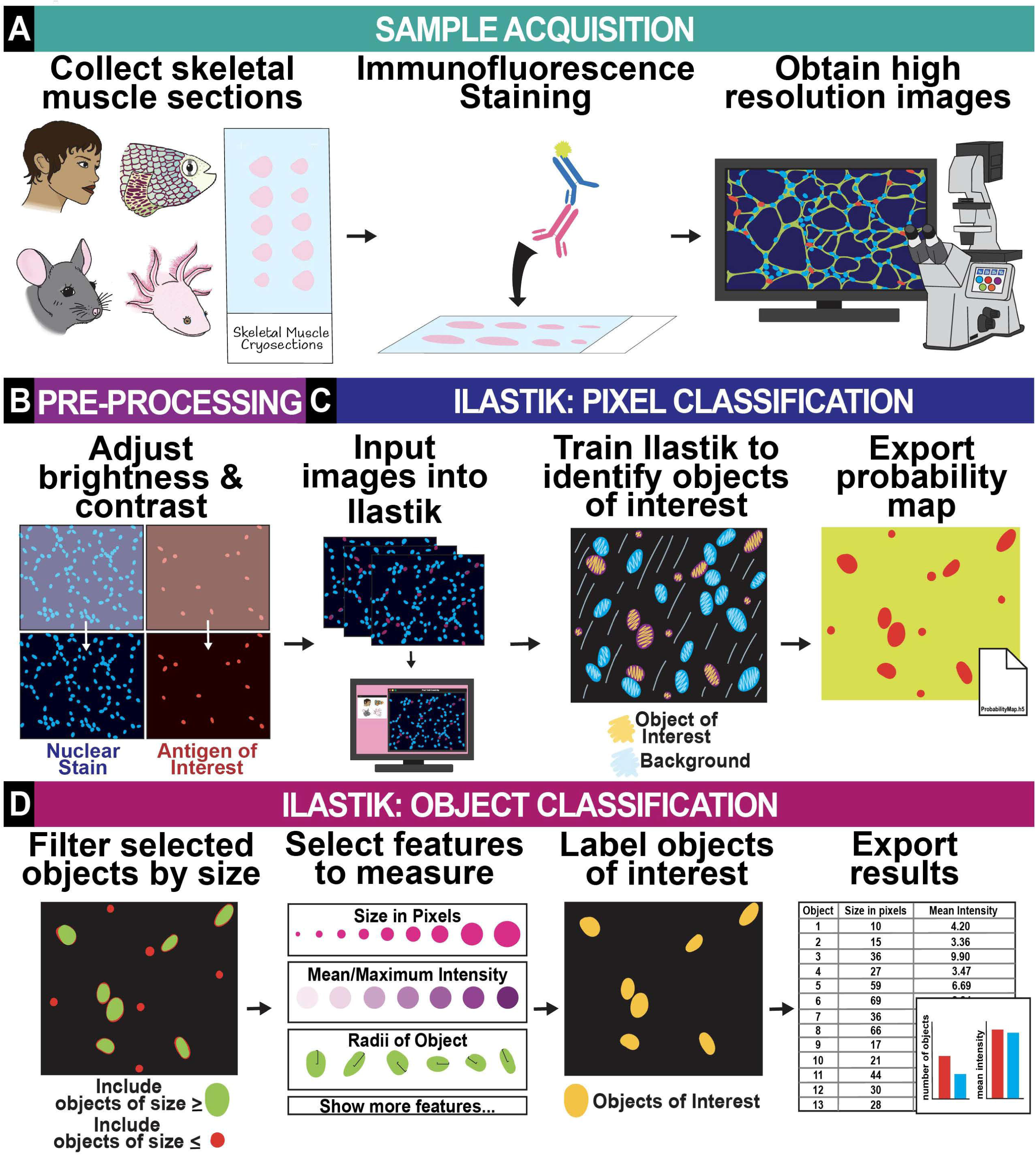
An overview of the Ilastik workflow. The Ilastik workflow can be broken down into four major steps: A) Sample acquisition; B) Pre-processing of images; C) Ilastik: Pixel Classification; and D) Ilastik: Object Classification. Briefly, (A) muscle sections are obtained from the desired model organism. Sections are then stained using IHC for the antigen of interest and Hoechst (nuclear) and then whole muscle sections are imaged using a microscope. (B) Brightness and contrast are adjusted to every image to ensure that both channels (i.e., nuclear stain and antigen of interest) have equal brightness. (C) Images are then loaded into Ilastik where Pixel Classification is used to make a training model, which is run on all images to generate a probability map demonstrating all objects of interest identified by the program. (D) In Object Classification, all objects are further filtered by size and additional features are chosen to report in final data (i.e., raw counts, size in pixels, mean/maximum intensity, etc.). The final selection of desired objects is easily graphed in Excel or Prism GraphPad.

To evaluate the broad utility of Ilastik in quantifying cell fates in multiple vertebrate species, we obtained skeletal muscle cryosections from multiple vertebrate model systems including mouse (*M. Musculus*), human (*H. Sapiens*), axolotl salamanders (*A. mexicanum*), and short-lived, African turquoise killifish (*N. furzeri*, *GRZ strain*). Next, we stained all the sections with a fluorescently labeled PAX7 antibody along with Hoechst to identify PAX7+ MuSCs. PAX7 is a bona-fide marker gene which is uniquely expressed in quiescent and proliferating MuSCs in mammals^15–17^; and whose expression is required for maintaining stem cell identity and mediating a full recovery of muscle after myotrauma^18–20^. We then obtained high resolution, 16-bit images of the entire muscle sections at 20X magnification, since we find that this is an adequate resolution to capture the fine details of the tissue that are critical for accurate quantification in Ilastik.

The pre-processing phase of the pipeline almost always requires adjustments to the brightness and contrast of the images to make sure the signal intensities between the antigen of interest (in our case, PAX7) and the nuclear stain (Hoechst) are as equivalent as possible.

Next, the pre-processed images are loaded into Ilastik to start Pixel Classification. In this phase, we teach Ilastik how to count PAX7+ cells. We achieve this by training the software to distinguish between objects of interest (i.e., PAX7+ cells) from objects we do not want included in our analysis such as non-PAX7+ cells in the tissue and non-specific staining (i.e., background). After user-specified training, Ilastik applies the same instruction to the entire tissue section and then generates a probability map that classifies each feature in the image as an object of interest (i.e., PAX7+ cells) or an object to exclude (i.e., non-PAX7+ cells, non-specific staining, unstained space). We note that training can be applied to several images at once with a feature in Ilastik called Batch Processing.

Finally, we input the pre-processed images and their accompanying probability map for the second portion of the analysis: Object Classification. In this step, we refine what Ilastik interprets as an object of interest by (1) establishing the level of congruency at which an object is labeled (which is important to faithfully capture the size of PAX7+ cells) and (2) by filtering selected objects by their size in pixels. Collectively, this workflow allows us to count objects of a specified size, which is useful if we are interested in only small objects (i.e., cells). Additionally, we can also capture other measurements of interest such as size in pixels, mean/maximum intensity, radius, and more. At the end of the analysis workflow, Ilastik generates a spreadsheet displaying the number of objects counted and their respective feature measurements. Remarkably, Ilastik can analyze an entire cohort of IHC-stained muscle sections in a matter of hours, whereas manually counting would take days to even weeks to complete the same task.

### Highlighting key steps in the Ilastik workflow

In this section, we will expand on three of the most crucial steps in the Ilastik pipeline that greatly impacts the accuracy of the results. The first major step is to ensure that each image has an optimal brightness and contrast (B/C) prior to analysis. Essentially, Ilastik requires a composite image displaying both PAX7 (green) and Hoechst (blue) immunofluorescent signals since it will only count cells that display both signals (**Figure 2A**). In an ideal situation, where PAX7 and Hoechst are of equal intensity, a merged image containing both channels will generate a cyan color (**Figure 2B**). To check if the merging is occurring appropriately, we recommend generating a Red, Green, Blue (RGB) histogram displaying the distribution of pixel intensities in the red, green, and blue channels. With an optimal B/C adjustment, most of the pixels display the brightest green and blue signals, which results in cyan-colored PAX7+ cells when both channels are merged (**Figure 2B**). On the other hand, in an inadequate B/C adjustment most pixels display very bright blue signal but weaker green signal, which results in fewer cyan-colored PAX7+ cells that will be confidently identified by Ilastik (**see Figure S1A**). In other words, if PAX7 signal is not equal to Hoechst, there is a higher probability that one will exclude out cells with lower expression levels of PAX7 given that these objects will appear blue and mask low levels of green.

**Figure 2.**
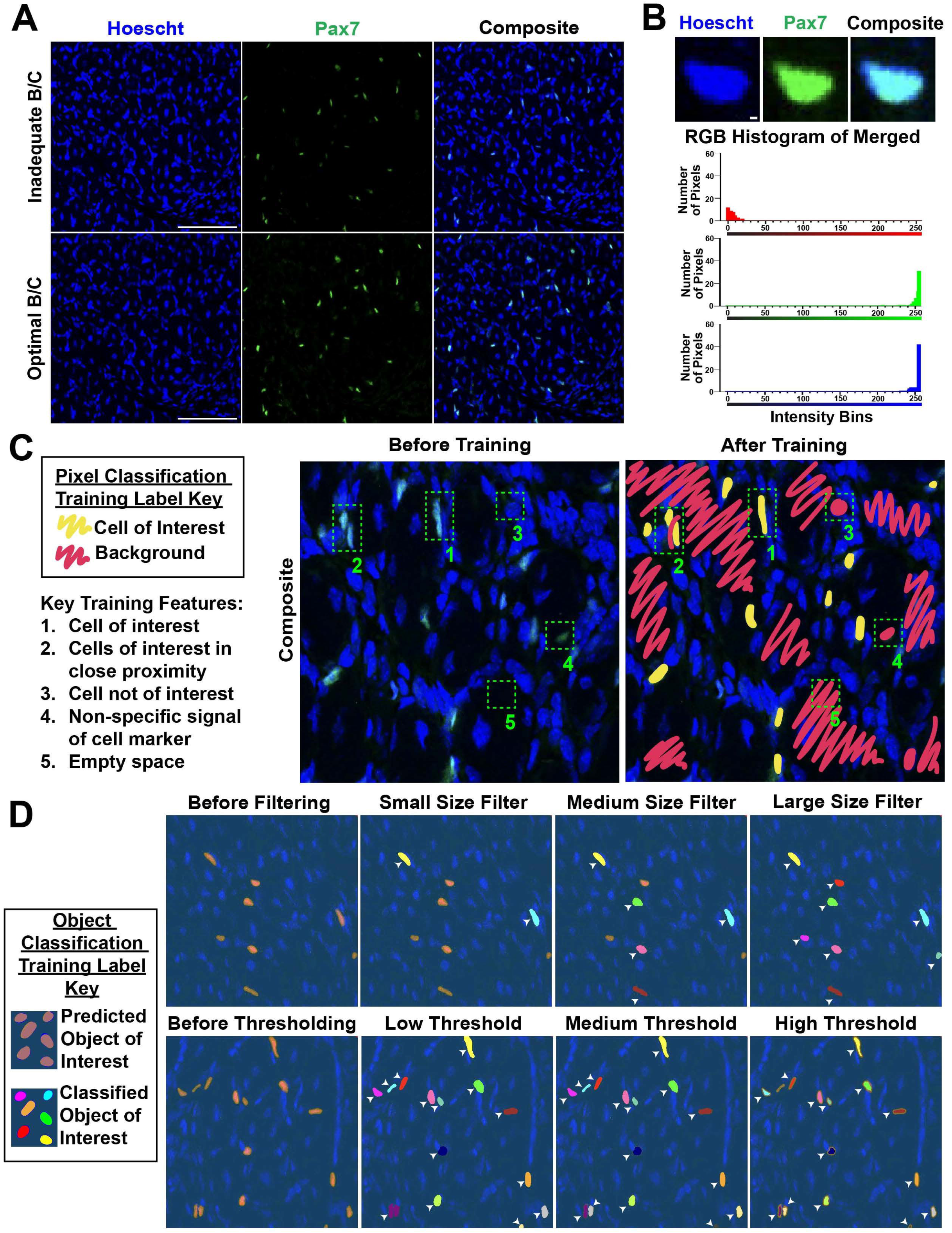
Highlighting key steps in the Ilastik workflow. (A) Representative 10X image of an inadequate adjustment to brightness/contrast (B/C) where green objects are dimmer than blue objects leading to fewer noticeable cyan PAX7+ cells in the composite image (top). Representative 10X image of an optimal adjustment to brightness/contrast where green objects are equal in brightness to blue objects resulting in more noticeable PAX7+ cells in the composite image (bottom). Scale bar represents 100μm. (B) Example of one cell displaying an optimal equal brightness of nuclear stain (Hoechst) and PAX7 (Green) (top). Representative histogram showing what an optimal brightness and contrast adjustment looks like when looking at pixel intensity of the red, green, and blue channels (bottom). Scale bar represents 1μm. (C) Legend showing our labeling scheme and several of the key training features appropriately marked in the representative images used for Pixel Classification in Ilastik. Images showing 5 boxes marking key features prior to training (left) and an example of what the image looks like after a successful training model has been applied (right). (D) Representative images of objects before (depicted with pale orange color) and after adjustments to filtering/thresholding. Objects that are positively selected during filtering/thresholding in Object Classification are marked with a variety of bright colors. Filtering by size allows one to separate small from large objects, and thresholding ensures objects are labeled congruently and not merging with neighboring objects of interest. White arrows in the image show how adjusting the size filter or thresholding (low, medium, and high) impact the ability to adequately identify all PAX7+ cells (brightly colored objects) versus objects that are not identified as PAX7+ cells (pale orange color).

The second critical step in the Ilastik pipeline is the process by which the investigator trains the program how to identify objects on interest on a set of images during Pixel Classification. Here, we highlight 5 key training features that we incorporate into our workflow to ensure consistent and reproducible analyses (**Figure 2C**). During the training phase, we make sure to (1) label 50-100 PAX7+ cells spanning at least 3 images as objects of interest, ensuring that we are training from multiple images that came from different slides or conditions. This is important as staining efficiencies can vary from one slide to another, and thus, these variations need to be captured in your training model. In addition, we choose examples from low, medium, and high expression levels for PAX7, as well as PAX7+ cells that range from round to more elongated in shape (**Figure 2C**). Next, when two or more PAX7+ cells are in close proximity (2), we carefully label the thin space separating them as background to avoid these cells from being recognized as one large cell. We also extensively label (3) PAX7-nuclei (blue signal only), (4) non-specific staining for PAX7 (green signal not overlapping Hoechst), and even (5) areas lacking any staining (i.e., empty black space) as background. Lastly, a useful feature of the training component is the Live Update function, which enables one to make quick adjustments (i.e. label additional objects or delete objects) and see in real time what Ilastik is classifying as objects of interest versus excluded objects based on the updated training model. After pixel classification is done a probability map identifying objects of interest is generated and exported for use in the following steps.

The next key step in the Ilastik workflow is Object Classification where one can finetune what one wants Ilastik to count as an object of interest using two parameters called (1) size filtering and (2) thresholding. After training, but before size filtering and thresholding, all objects are labeled in a pale-orange color representing the probability map generated from Pixel Classification (**Figure 2D**). Once filtering and thresholding of objects is completed, newly selected objects based on the training model are labeled in bright colors. The size filter selects objects by their size in pixels, where small-range filters will exclude many objects while large-range filters will capture most if not all objects (**Figure 2D**). Thresholding labels objects based on the probability of each pixel in the object having all the selected features of interest, which essentially dictates how pronounced the object will be labeled. A low threshold will lead to a label that is larger than the object itself due to the inclusion of low probability regions around the object of interest. However, as the threshold is increased, the objects will be marked with smaller labels that are more consistent with their natural shape (**Figure 2D**). Both parameters are applied simultaneously, so it is important to set a size filter and threshold that will enable you to select all the desired cells. Object Classification also includes a Live Update feature which allows one to visualize and adjust parameters in real time. Finally, it is important to note that the size filter and threshold parameters will likely vary in every experiment even if one is quantifying the same cell type due to changes in size/shape in different conditions (e.g. uninjured, injured, aged, young, and vary from species to species).

### Ilastik accurately quantifies PAX7+ cells in mouse skeletal muscle tissue before and after myo-injury

To demonstrate Ilastik’s accuracy in automatically quantifying PAX7+ cells in skeletal muscle, we performed PAX7 IHC in uninjured and injured skeletal muscle sections followed by quantification of PAX7+ cells using (1) Ilastik or (2) by 3 trained manual counters in our laboratory. Specifically, we injured the tibialis anterior (TA) muscles of C57BL6/J mice with 1.2% Barium Chloride (BaCl_2_) and then harvested the muscles 7 days post-injury (dpi) (**Figure 3A**). Interestingly, we found that the number of PAX7+ cells in whole TA muscle sections detected by Ilastik and 3 manual counters was essentially the same when evaluated separately in both conditions (i.e., no injury versus 7dpi) (**Figure 3B-C**). More importantly, we see a similar ∼6-fold increase in PAX7+ cells in skeletal muscle 7dpi when compared to uninjured muscle using Ilastik and manual counting (**Figure 3D**), similar to trends reported in previous studies^21,22^. These data suggest that Ilastik accurately quantifies PAX7+ cells in skeletal muscle before and after injury and preserves biological trends to a level and rigor that is comparable to manual quantification in a fraction of the time (i.e., hours versus multiple days).

**Figure 3.**
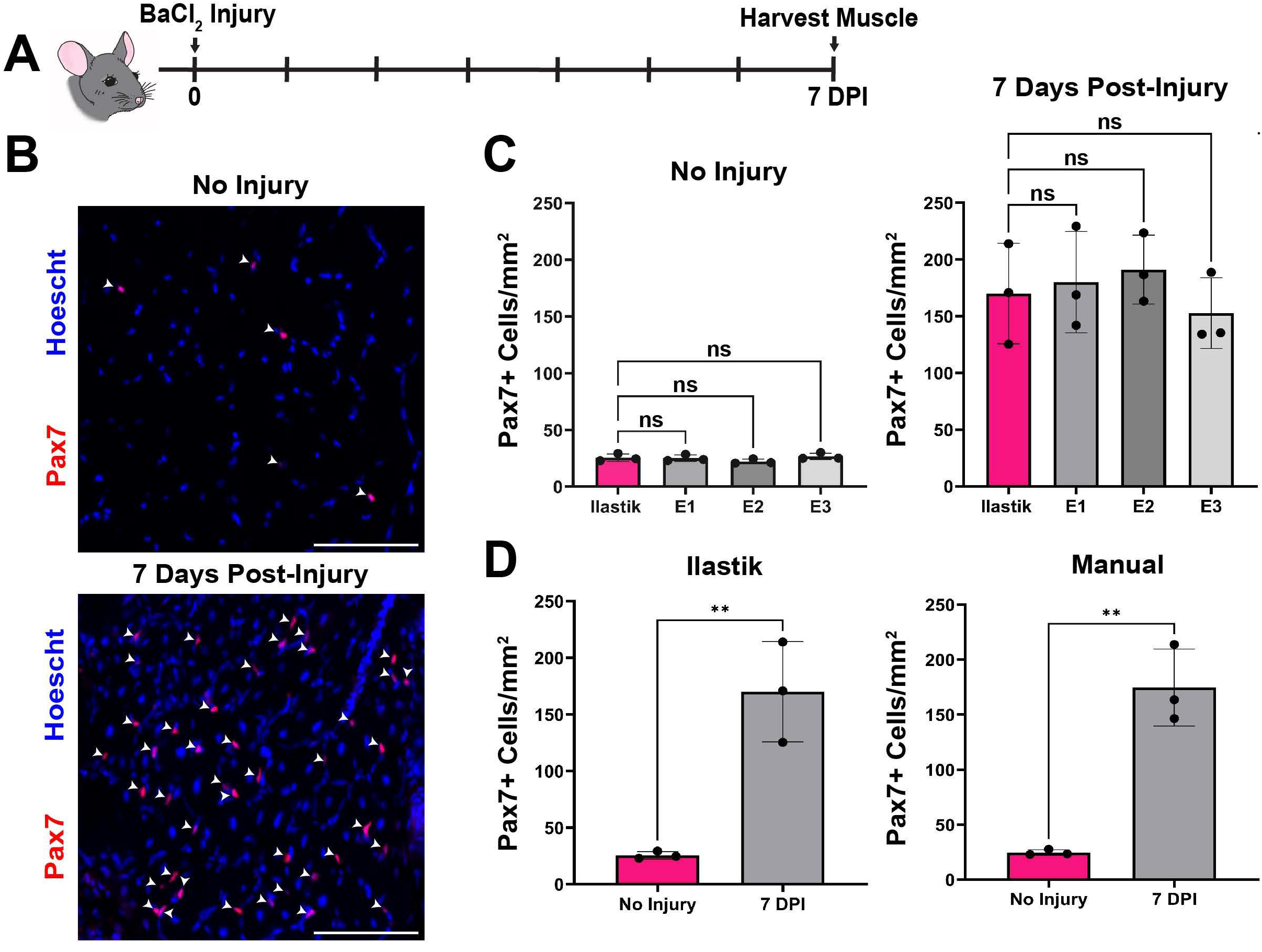
Ilastik accurately quantifies PAX7+ cells in mouse skeletal muscle tissue before and after myo-injury. (A) Schematic showing that Tibialis Anterior (TA) muscles were injured with Barium Chloride (BaCl_2_) and then harvested for analysis after 7 days post injury (dpi). (B) Representative 20X image of mouse TA with no injury and 7 dpi immuno-stained for PAX7 (red) and Hoechst (blue). White arrows denote Pax7+ cells. (C) Bar plot showing PAX7+ cells/mm^2^ calculated by Ilastik and manually by 3 trained counters in the laboratory of uninjured TAs (Left) and 7dpi TAs (Right) (N = 3 depths per time-point). No statistical difference between Ilastik and manual counters. (D) Bar plots comparing PAX7+ cells/mm^2^ in TA muscle with no injury versus 7dpi as quantified by Ilastik (Left) and manually by 3 trained manual counters (Right). Statistical comparisons between population means were made using a (C) one-way ANOVA and a (D) unpaired, two-tailed, Student’s t-test with a p-value cutoff of 0.05 for significance. The difference in PAX7+ cells/mm^2^ between conditions (in both Ilastik and manual counting) was found to be statistically significant (** pσ;0.01). NS means not significant in our statistical test. Scale bars represent 50μm.

### Ilastik precisely quantifies PAX7+ cells in human muscle samples

To test whether our Ilastik analysis pipeline can also identify PAX7+ cells in human skeletal muscle, we first obtained human muscle biopsies from the subscapularis and deltoid muscles of 2 male patients (28 yrs, 56 yrs) (**Figure 4A**). After freezing the tissues, sectioning, and performing IHC for PAX7, we took images of the entire muscle section for the two samples. Consistent with our analysis using mouse tissues, we found that Ilastik accurately identified PAX7+ cells throughout the skeletal muscle section (**Figure 4B-C**). Importantly, when we quantified the abundance of PAX7+ cells within each muscle section, we found that Ilastik was able to quantify PAX7+ cells to a level comparable to that of 3 manual counters in both muscle samples (**Figure 4D**). Together, these data show that Ilastik can speed up the downstream analysis of counting PAX7+ cells in clinically relevant human muscle biopsy samples.

**Figure 4.**
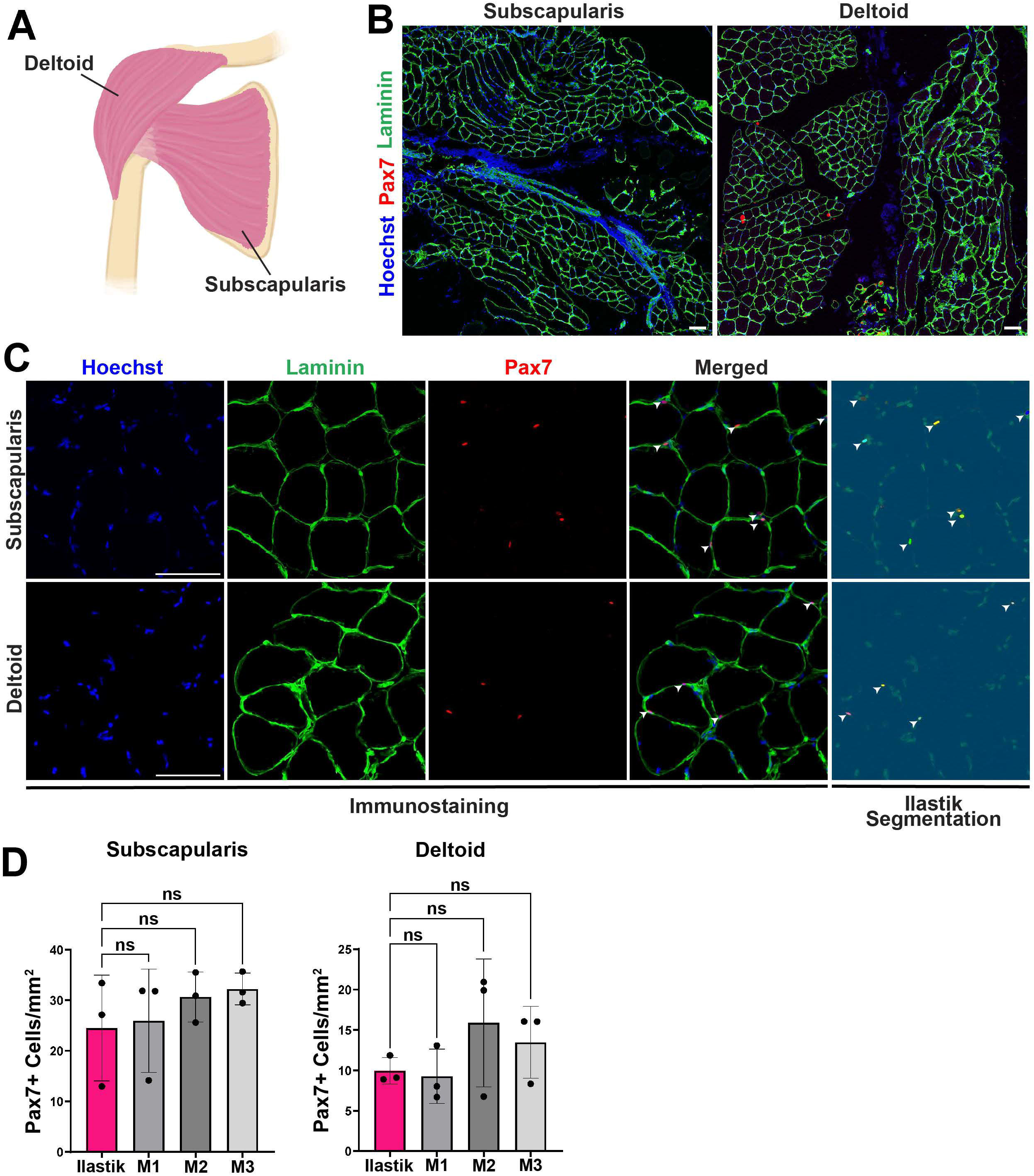
Ilastik precisely quantifies PAX7+ cells in human muscle samples. (A) Schematic showing the location of each skeletal muscle biopsy from 2 human patients: 1) subscapularis (28yo, male); and 2) Deltoid (56yo, male) muscles. (B) Representative 10X images of subscapularis and deltoid muscle immuno-stained for PAX7 (red), Laminin (green), and Hoechst (blue). (C)20X representative images showing PAX7 (red) co-localized with Hoechst (blue). All PAX7+ cells are marked by white arrowheads. (D) Bar plots showing the number of PAX7+ cells /mm^2^ in subscapularis (Left) and deltoid (Right) muscles calculated by Ilastik and 3 manual counters. Data are expressed as mean +- SD (N=3 depths per human muscle sample). Statistical comparisons between population means were made using a one-way ANOVA with a p-value cutoff of 0.05 for significance. NS means not significant in our statistical test. No significant difference was observed between Ilastik vs trained cell counts. Scale bars represent 100μm.

### Ilastik efficiently measures PAX7+ cells before and after tail amputation in axolotl salamanders

To further evaluate the broader use of Ilastik in quantifying IHC-stained tissue sections in a non-mammalian vertebrate species, we decided to challenge Ilastik with identifying PAX7+ cells in the super-healing, axolotl salamander (*Ambystoma mexicanum*). Axolotl salamanders are a highly regenerative species that regenerates whole appendages including arms, legs, and tails after amputation^23–28^. It is generally thought that following amputation, the regenerating appendage forms a unique structure called a blastema, which contains all the stem cells and progenitors needed to rebuild a complex tissue including skeletal muscle^23,29^. Thus, we isolated tails from axolotl salamanders before and 14 days post amputation (dpa), marking the beginning of the blastema stage, and then quantified PAX7+ cells at both time-points using Ilastik or manually by 3 trained counters (N=3 tails per time-point with 3 depths averaged per tail) (**Figure 5-C**). Consistently, we found that Ilastik and manual counting yielded a similar amount of PAX7+ cells in tail skeletal muscle before and 14 dpa (**Figure 5D-E**). Additionally, our data confirms that pre-existing PAX7+ cells occupy tail skeletal muscle prior to injury in axolotl salamanders and, more interestingly, that they are present in the blastema of the regenerating tail ^30,31^. Altogether, we show that Ilastik is very efficient at quantifying PAX7+ cells in whole appendages in a super-healing species.

**Figure 5.**
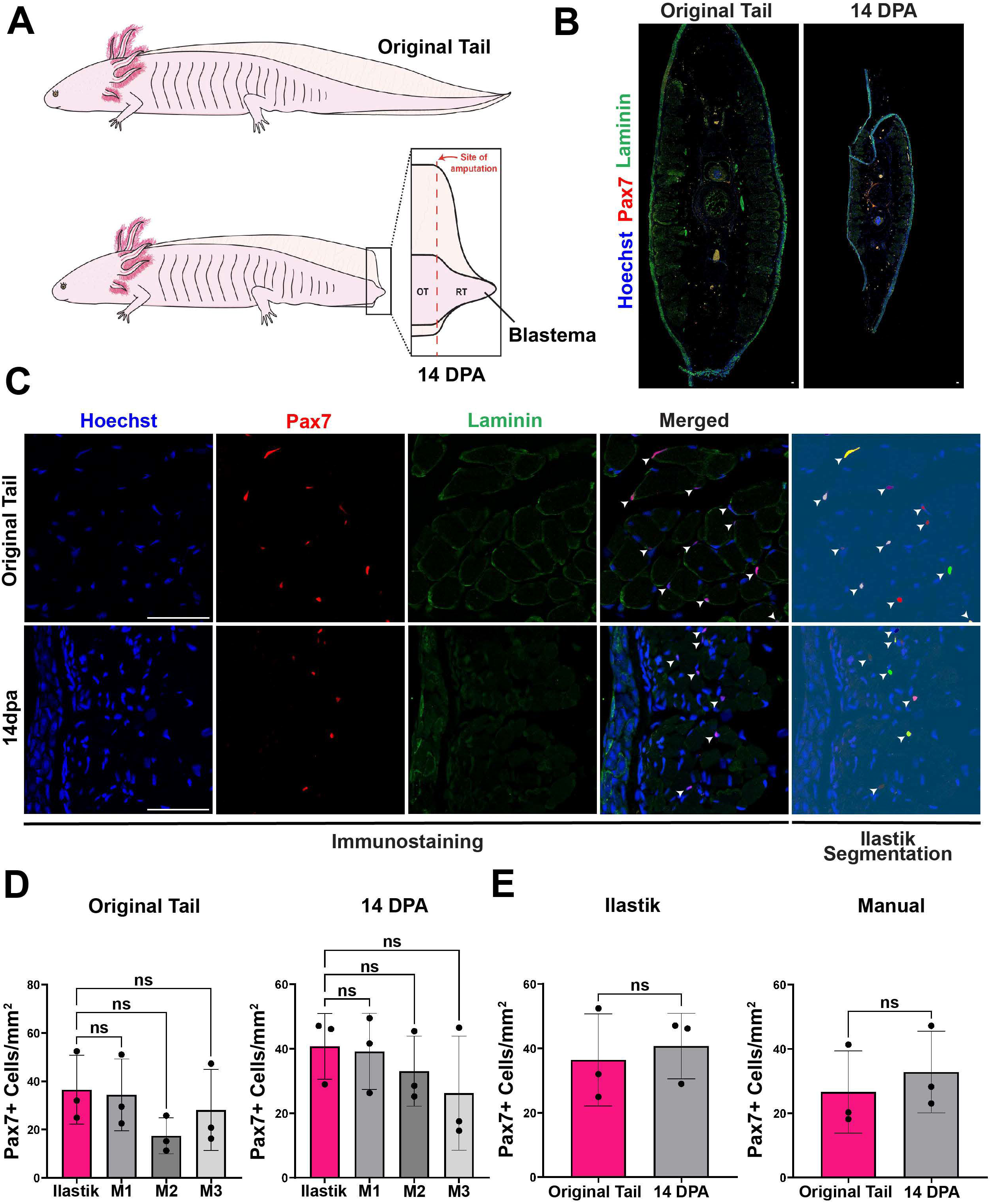
Ilastik efficiently measures PAX7+ cells before and after tail amputation in axolotl salamanders. (A) Schematic showing our experimental plan: salamander tails were harvested before and 14 days after tail amputation (i.e., the blastema phase). (B) Representative stitched 20X image of a transverse section of an axolotl salamander tail before and 14 days post tail amputation showing PAX7 (red), Laminin (green), and Hoechst (blue). (C) Zoom in representative 20X images showing colocalization of PAX7 (red) and Hoechst (blue) marked by arrowheads. Ilastik segmentation shows all PAX7+ cells as objects that are detected using Ilastik after an appropriate size and threshold cutoff. (D) Bar plots showing the number of PAX7+ cells/mm2 in the original (uninjured) and 14 days after tail amputation calculated by Ilastik and 3 manual counters. (E) PAX7+ cells/mm^2^ before and 14 dpa in axolotl salamanders measured by Ilastik (Left) and 3 manual counters (Right) separately. Statistical comparisons between population means were made using a (D) one-way ANOVA and a (E) unpaired, two-tailed Student’s t-test with a p-value cutoff of 0.05 for significance. Data are shown as mean +- SD (N=3 tails per time-point, with 3 depths averaged per tail). NS means not significant in our statistical test. Scale bars represent 100μm.

### Ilastik measures PAX7+ cells in whole trunks of young and old African turquoise killifish

We next decided to test whether Ilastik could accurately identify PAX7+ cells in an emerging naturally short-lived teleost fish model, the African turquoise killifish (*Nothobranchius furzeri*) (**Figure 6A**). The turquoise killifish is best known for being the shortest-lived vertebrate species that are bred in captivity, potentially making them a more relevant model for studying vertebrate aging^32–34^. Therefore, we decided to study a strain of turquoise killifish (GRZ) that reaches sexual maturity at 4 weeks of age and dies of old age by 4-6 months^35,36^. Although a longer-lived MZCS_08/122 strain of the African Killifish (living approximately 1 year) has been recently established as a new model of muscle aging^37^, whether the even shorter-lived GRZ strain, which dies within 4-6 months^38^, shows a more rapid muscle aging phenotype has yet to be determined.

**Figure 6.**
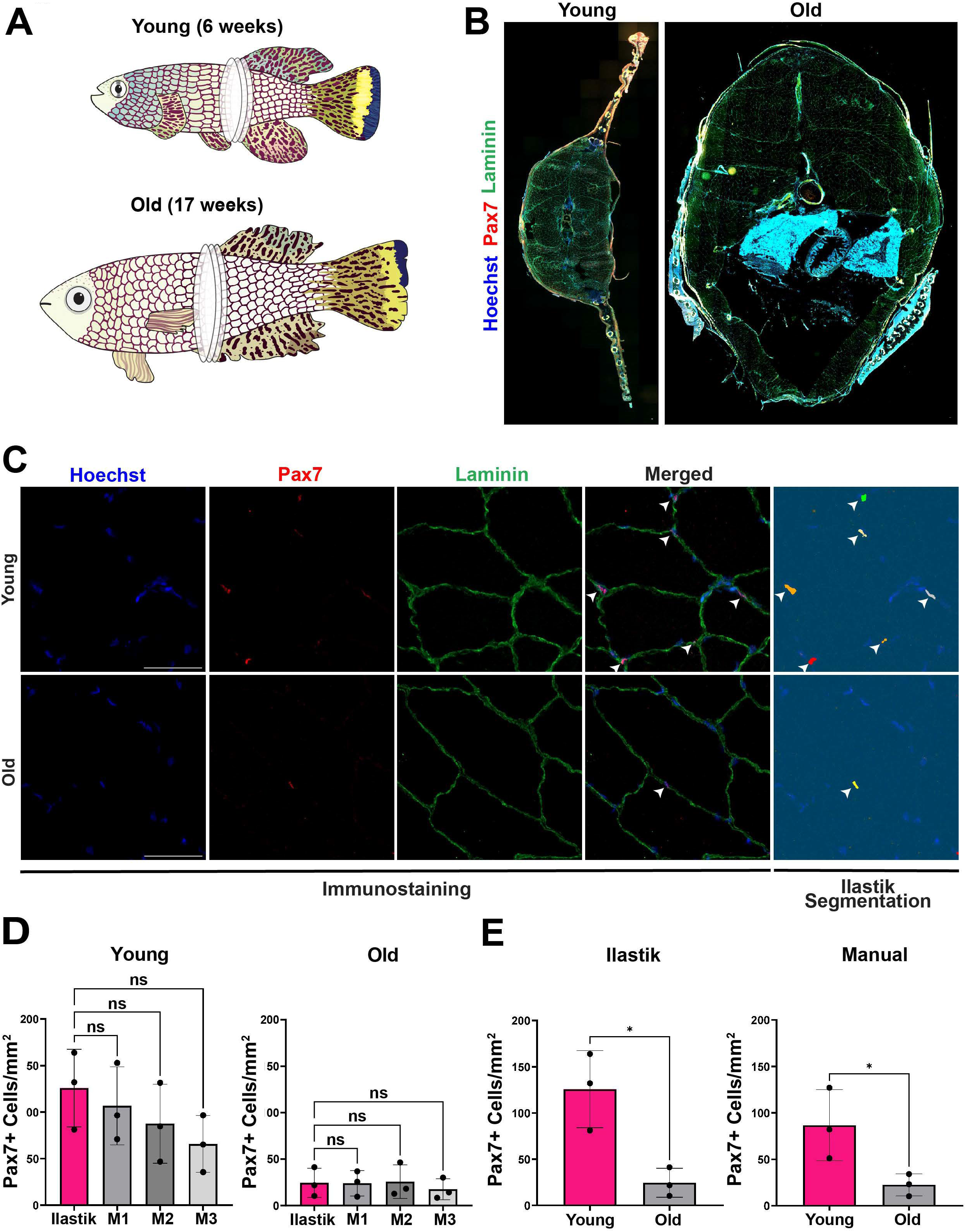
Ilastik measures PAX7+ cells in whole trunks of short-lived young and old African Turquoise Killifish. (A) Schematic showing young (6 weeks) and old (17 weeks) turquoise killifish and the trunk region with 3 different depths that were used for immunostaining. (B) Representative stitched 20X image of transverse sections from young and old trunk region, PAX7 (red), Laminin (green), and Hoechst (blue). (C) Zoom in 40X stitched images showing colocalization of PAX7 (red) and Hoechst (blue) marked by arrowheads. Ilastik segmentation shows all PAX7+ cells as objects that are detected using Ilastik after an appropriate size and threshold cutoff. (D) Bar plots showing the number of PAX7+ cells/mm^2^ in young (Left) and old (Right) turquoise killifish trunks measured by Ilastik and by manual counters. (E) Bar plots showing PAX7+ cells/mm^2^ between young and old trunks measured by Ilastik (Left) and 3 manual counters (Right) separately. Essentially, Ilastik and manual counts show a comparable statistically significant trend towards lower PAX7+ cell numbers in old relative to young turquoise killifish. Statistical comparisons between population means were made using a (D) one-way ANOVA and (E) an unpaired, two-tailed, Student’s t-test with a p-value cutoff of 0.05 for significance(*=p<0.05). Data are shown as mean +- SD (N=3 fish per condition (young/old) with 3 depths averaged per fish). Scale bars represent 50μm.

Thus, we obtained young (6 week) and old (17 week) male GRZ turquoise killifish and harvested, cryo-preserved, and sectioned the trunks (i.e., containing large muscle groups) (**Figure 6A**). Subsequently, we performed IHC for PAX7 on young and old trunk sections and then imaged the entire sections (N= 3 fish for young and old with 3 depths averaged per fish) (**Figure 6B-C**). Similar to the performance in mice, humans, and axolotl salamanders, Ilastik accurately identified PAX7+ cells to a level that was similar to 3 manual counters in both young and old fish (**Figure 6D**). It is worth noting that the variability between the 3 manual counters was greatest in the turquoise killifish when compared to mouse, human and axolotl salamanders, which we suspect is due to user error when having to manually count a whole fish trunk (very large samples that took approximately 2 hours per section). Most importantly, we found that adult (6-week-old) male turquoise killifish have a pre-existing PAX7+ cell population throughout the trunk that is dramatically reduced with age (17-week-old) in the GRZ strain when quantified by Ilastik and by 3 manual counters (**Figure 6E**). Overall, these data highlight the power of Ilastik to quantify PAX7+ cells in whole trunks to a level that exceeds human capabilities, and thereby, allowing us to uncover a decline in PAX7+ cells in aged trunks of the extremely shorted-lived, turquoise killifish (GRZ strain).

## Discussion

One of the most challenging aspects of studying vertebrate tissue and organ regeneration in the laboratory is the time it takes to analyze many multi-colored, IHC-stained images—capturing changes in stem cell fates after tissue injury, after drug-treatment, or in pathogenic states. Historically, this type of analysis has been dominated by manual quantification in a small subset of randomly selected images from the larger tissue sample. However, in an ideal situation, an investigator would analyze the entire tissue section and generate a conclusion based on the entire data and not just a small subset of the data. Thus, new technologies to facilitate the automatic analysis of IHC-stained images representing entire tissues sections are needed to remove human bias and analyze the resulting data in a more consistent and reproducible way. In this study, we benchmarked a newly developed ML-based imaging analysis platform called Ilastik^14^ and demonstrated that it can quickly and accurately quantify PAX7+ cells before and after myo-injury in multiple vertebrate species to levels that are comparable to manual quantifications performed by trained investigators. Our success in optimizing Ilastik for studying vertebrate muscle biology suggests that the Ilastik pipeline could be adapted to other antigens and tissue/organ types.

Indeed, there have been several programs previously developed with the intent of replacing manual quantification of IHC stained images of skeletal muscle, especially for automatically quantifying PAX7+ cells and muscle fiber sizes^39–43^. Regarding the quantification of PAX7+ cells in muscle sections, there is only one other free program called MuscleJ^39,40^, which is an add-on macro that can be used in ImageJ (Fiji). However, a major limitation in using MuscleJ is that PAX7+ cells are defined as those co-associated with laminin-positive myofibers. Therefore, this absolute requirement makes it difficult to detect PAX7+ cells when they co-associate with damaged or small regenerating myofibers displaying no to weak laminin expression, which is typical within the first week after muscle injury *in vivo*. Given that Ilastik only requires co-association with a nuclear stain (i.e. Hoechst and DAPI), it is capable of quantifying PAX7+ cells alone or in association with newly formed myofibers very effectively within the first week after myo-injury (**Figure 3**), making this program the most versatile free resource currently available for the community.

One of the most striking aspects of our evaluation of Ilastik on whole tissue sections was generally how similar its values were when compared to manual quantification from several trained investigators in our laboratory. We anticipate that the speed and accuracy of Ilastik will increase the rigor by which we measure PAX7+ cells moving forward: by making it more feasible to quantify the entire muscle section and in a greater number of depths from the muscle. For example, it is the gold standard to quantify multiple depths (∼3-6) from an uninjured, control or experimentally perturbed muscle (i.e., TA, quadriceps muscles, etc.,). To do this, one must image a cross section of the entire muscle from a minimum of 3 depths per muscle. Depending on the effect size of your phenotype, you would need 3-10 biological replicates, meaning anywhere from 9 to 30 individual sections to analyze if you had only 2 groups (control and experimental). Obviously, with more than 2 groups or a time-course, the number of whole sections to quantify may double or even triple. This would be very labor intensive and likely prone to mistakes due to fatigue that an investigator would experience when manually quantifying an entire tissue section, which is exactly why many continue to quantify representative images from each muscle section. Consistently, we noticed that as the area to be manually counted increased (particularly in axolotl salamanders and turquoise killifish) there was more variability between counts for trained investigators that are quantifying the same images. Instead, after successfully creating a training model on multiple images, all images are simply uploaded to the program and Ilastik quantifies PAX7+ cells with a remarkable accuracy.

Another benefit of using the Ilastik workflow is that it encourages fields that were previously more qualitative to take a more quantitative approach. For example, our laboratory recently started studying MuSC activation in multiple vertebrate species including non-traditional model organisms like axolotl salamanders and African turquoise killifish. These super-healing species are interesting because they can rebuild muscle *de novo* after trauma in fins, arms, legs, and even tails^33,29^. In our initial studies of detecting PAX7+ cells in these super-regenerating species, we ran into a roadblock when trying to perform manual quantifications in the same way we had previously done in mouse TA muscles. While it takes a considerable amount of time to quantify PAX7+ cells in whole mouse TA sections manually, it takes even longer to do the same manual counts in whole cross sections of the trunk for fish or the appendage of a salamander, due to the increase in size of the sections being analyzed. Additionally, if one wants to apply the same rigor of quantifying 3-6 depths per single trunk, tail, or limb, and considering multiple biological replicates and/or an amputation time-course, it is completely impractical to do this using manual quantification. Thus, in our studies, we provide a new solution where we can accurately quantify PAX7+ cells in whole salamander tails before and after tail amputation, as well as in whole trunks from young and aged fish. Using this strategy, the field is now poised to study MuSC fate decisions in complex appendages with unprecedented rigor.

Bolstered by the speed and accuracy of Ilastik, we were able to make two additional biological observations in these non-traditional model organisms. First, in the axolotl salamander system, we found that similar to mammalian MuSCs^44^, (1) PAX7+ cells reside between skeletal muscle myofibers in the tail (**Figure 4C**,^30,31^); and that (2) PAX7+ cells were observed in the “regenerating” blastema 14 days after tail amputation, suggesting that MuSCs may be recruited into the blastema from adjacent uninjured muscle tissue to facilitate *de novo* muscle formation in later weeks. Consistent with this model, lineage tracing showed that a pre-existing PAX7+ cell population contributes to new muscle formation in the regenerating limb after amputation in vivo^30^. While the mechanism controlling the recruitment of axolotl salamander MuSCs into the blastema remains a mystery, we now have a way to quantify their movement in a rigorous and quantitative way throughout the regenerative process. Second, our studies using the short-lived, African turquoise killifish (GRZ strain)—which reaches adulthood at 1 month but has a captive lifespan of 4-6 months—showed a reduced number of PAX7+ cells per mm^2^ in aged (17 weeks) trunks relative to young (6 weeks) trunks. These data may suggest that as the fish grows in size during the aging process (**Figure 5B**), there may be less PAX7+ cells to cover a larger area in older trunk tissues, which presumably, would have negative consequences in the event of muscle trauma. We note that a recent study using a longer-lived turquoise killifish strain (MZCS_08/122) showed no change in PAX7+ cell numbers per myofiber between 6 weeks and 22 weeks, however they did observe a 2-fold decrease by 52 weeks (i.e., 1 year)^37^. Together, these data clearly suggest that the GRZ strain of the turquoise killifish shows promise as a model system for accelerated muscle aging in a vertebrate animal.

Although there are many benefits of using the Ilastik workflow, it is worth noting its caveats and limitations. First, we cannot stress enough that Ilastik is only as accurate as the investigator who generates the training model. Therefore, semi-automatic programs like Ilastik do not eliminate the need for investigator input, but rather, it relies heavily on well-trained users to teach them how to identify a positive cell. Second, the accuracy of Ilastik is very dependent on having high-quality IHC images, thus it is critical to optimize staining conditions before even attempting to use Ilastik. Third, because Ilastik requires a merged file to quantify a multi-colored object, we have found that it does well for 2- and 3-colored images. However, using more than 3 channels presents a challenge given it increases the chances that the desired objects in the merged image will approach a color similar to one of the original colors or background signal (i.e., shades of black or grey), thus, making the analysis erroneous.

Altogether, we highlight the benefits of Ilastik in expediting one of the most time-consuming manual measurements in MuSC biology (i.e., PAX7+ cell counting) in not just mice, but in 3 other vertebrate species including humans, axolotl salamanders, and fish. We are confident that a similar strategy discussed here can be adapted to detect other marker genes and in other tissue types, and thus, transforming how investigators analyze the many IHC-stained images resulting from studies in vertebrate animal biology.

## Acknowledgements

We thank Dr. Seth Ruffins in the USC Stem Cell Optical Imaging Facility for his help with operating the Zeiss AxioScan Slide Scanner. This work was supported by a T32 Training Fellowship in Development, Stem Cells, and Regenerative Medicine (2T32HD060549-11) and Howard Hughes Medical Institute Gilliam Fellowship to AZM; NIGMS R35 GM142395 to BAB; NIH R01 GM115444 to TPL; and Junior Faculty Awards from the American Federation of Aging Research (AFAR) and Baxter Foundation and an NIH R01 AR080753 to AEA.

## Author Contributions

AZM, KS, and AEA conceived the study. AZM performed all mouse studies and related analysis. KS performed experiments and data analysis for human, salamander, and killifish studies. AL assisted KS in obtaining human muscle samples from our clinical collaborators. FP provided human muscle samples for studies. TL provided salamander tails (before and after amputation) for use in our studies. AA and BAB provided young and old turquoise killifish samples for our studies. KS, AZM, AEA, VL and AI performed manual quantification of PAX7+ cells. KK assisted in the comparison between manual counting and Ilastik in an earlier iteration of this work, providing the premise for current figures. AZM created all illustrations depicted in this study. AZM, KS, and AEA wrote the paper. All authors edited and approved the final version of the manuscript.

**Supplemental information titles and legends**

## STAR★Methods

### Key resources table

The key resources table (KRT) serves to highlight materials and resources essential to reproduce results presented in the manuscript. The items in the table must also be reported alongside the description of their use in the method details section. Literature cited within the KRT must be included in the references list. Please do not add custom headings or subheadings to the KRT. We highly recommend using RRIDs (see https://scicrunch.org/resources) as the identifier for antibodies and model organisms in the KRT. To create the KRT, please use the template below or the [KRT webform](https://star-methods.com/). See the more detailed [Word table template](https://www.cell.com/pb-assets/journals/research/cell/methods/table-template1.docx) document for examples of how to list items.

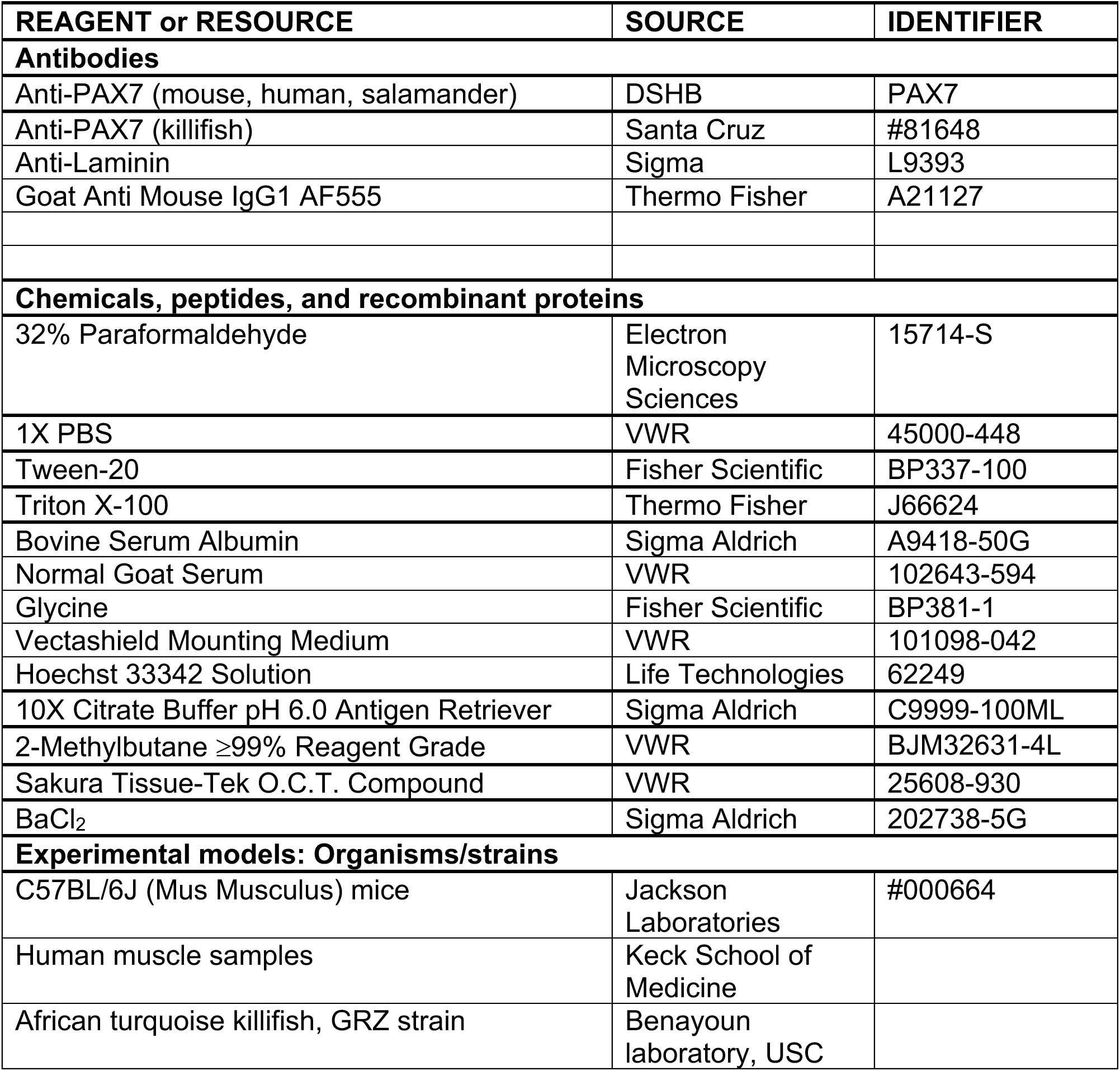

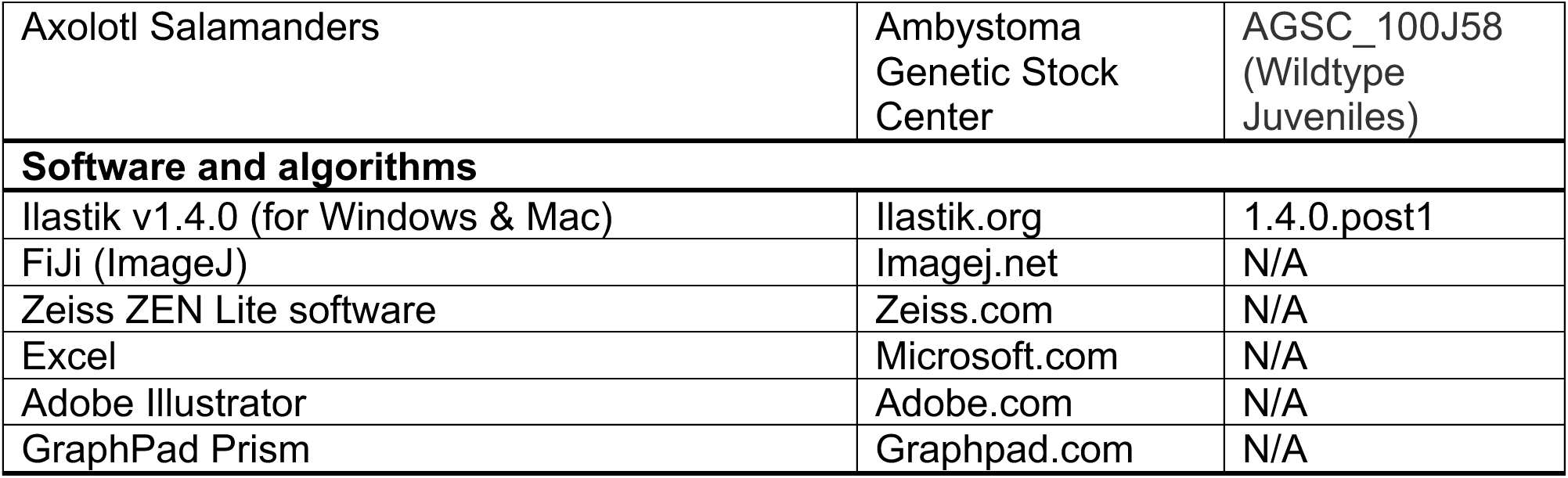

### Experimental model and study participant details

#### Mouse (Mus Musculus)

3-month-old male C57BL/6J (000664) mice were purchased from Jackson Labs and housed in our facility 2-6 weeks before the start of experimentation. All mice were housed in a 12-hour light/12-hour dark cycle and fed ad libitum and used according to our approved Institutional Animal Care and Use Committee (IACUC) protocol at the University of Southern California (USC) (Protocol: 21247).

#### Human (Homo Sapiens)

Human samples were obtained from patients undergoing surgery for shoulder injuries. Approximately 5mm of healthy muscle was removed from two male patients: 1) subscapularis muscle from a 28-year-old; and 2) deltoid muscle from a 56 year old. Informed consent was obtained before surgery and all procedures were performed according to our approved Institutional Review Board (IRB) protocol (HS-20-00506).

#### Salamander (Axolotl Mexicanus)

All axolotl salamander studies were performed according to the guidelines of the IACUC at USC (protocol 20992). Wild-type axolotl salamanders (5-8 cm snout-vent-length juveniles) were obtained from the Ambystoma Genetic Stock Center. Axolotl salamanders were housed individually in plastic aquaria at water temperatures of 18–20 °C under 12-h light/12-h dark schedules and fed a diet of salmon pellets 5 days per week. For tail amputation surgeries, axolotl salamanders were anesthetized by placing animals in baths of 0.03% benzocaine solution, and sterile scalpel blades were used to amputate tails hallway along original lengths. Following 14 days of regrowth (Blastema stage), regenerated tail samples were collected via second amputations 1 cm proximal to original amputation planes.

#### African turquoise killifish

African turquoise killifish (GRZ) strain was grown following standard accepted procedures in the field. Briefly, to limit aggression, all adult fish were singly housed in 2.8L tanks, on a recirculating aquatics system manufactured by Aquaneering Inc. System. System parameters were kept constant as follows: temperature: 29°C; pH: 7.3; conductance: 670-750 μS; ammonia and nitrite: 0ppm; Nitrate: ∼15 ppm. Adult fish were fed twice per day with Hikari Freeze Dried Bloodworms 4-6 hours apart, and live Artemia once per day. Fry (baby fish) were reared in system water incubated at 28°C, then placed on the recirculating system starting at 2 weeks after hatching. Fry were fed live Artemia exclusively until 4 weeks after hatching. The fish room was kept on a light/dark cycle of 13/11 hours (lights on 9am-10pm). The fish were euthanized by immersion in 1 g/L of Tricaine MS-222 dissolved in system water, followed by decapitation as a secondary method of euthanasia. Fish were always euthanized between 10 am and 12 pm to minimize circadian effects. The University of Southern California IACUC approved all husbandry conditions and experimental procedures under approved protocol 20879 for *Nothobranchius furzeri*.

#### Method details

##### Histology and preparation of muscle tissue samples

For mouse studies, the left TA muscle was injured via injection of 35ul 1.2% BaCl_2_ and the right TA was left uninjured. Both TAs were harvested 7 days post-injury, embedded in O.C.T. compound, and then cryo-preserved with 2-methylbutane cooled with liquid nitrogen for 45 seconds, transferred to dry ice, then stored at −80°C until needed. 12μm cryosections were collected from the entire length of the TAs using the Leica CM 1860 UV cryostat and stored at −80°C until ready for IHC.

For human samples, muscle biopsies were collected from healthy muscle tissue during surgery. The muscle samples were collected in 30% sucrose in PBS. The freshly collected biopsies were embedded in O.C.T compound followed by snap freezing in 2-methylbutane cooled with liquid nitrogen. 12μm serial cryosections were collected using a Leica CM 1860 UV cryostat and stored at −80°C until ready for IHC.

For Salamander, we collected samples from 10-13month old female Axolotl. Tails were amputated 1cm from the posterior vent of the animals. Tail samples were collected from uninjured (original tails) or 14 days post amputation (14dpa) animals and fixed in 4%PFA overnight at 4°C followed by several washes with PBS. Samples were then embedded in O.C.T compound and frozen in 2-methybutane cooled with dry ice. 16μm serial cryosections were collected using a Leica CM 1860 UV cryostat and stored at −80°C until ready for IHC.

African turquoise killifish were collected at different ages, 6 weeks for young and 17 weeks for aged fish. Only males were used for this study. Whole fish bodies were frozen in cryomolds in O.C.T compound by immersing in 2-methylbutane cooled with liquid nitrogen. The frozen cryomolds were stored at −80°C until sectioned. 12μm serial cryosections were collected using a Leica CM 1860 UV cryostat and stored at −80°C until ready for IHC.

##### Immunohistochemistry (IHC)

For mouse IHC of TA muscles, slides were fixed with 4% PFA for 5 minutes at room temperature (RT). Slides were washed with PBST (0.1% Tween-20 in PBS) for 5 minutes at RT 3 times (washes repeated after every step). Slides were incubated in 0.1M glycine for 5 minutes at RT. Antigen retrieval was performed on slides as follows: (1) slides were placed in a vertical slide rack and submerged in 1X citrate buffer (10X citrate buffer diluted in double-distilled water); (2) citrate buffer was brought to a boil 6 times using a microwave; and (3) slides were then washed with double-distilled water for 5 minutes at RT 3 times. Slides were permeabilized with 0.3% Triton X-100 for 20 minutes at RT. Slides were then blocked with blocking buffer (3% bovine serum albumin, 5% normal goat serum, 0.1% Triton X-100) for 1 hour at RT. Slides were incubated with primary antibody solution (1:2 DSHB mouse anti-PAX7 diluted in blocking buffer) overnight at 4°C. Slides were then washed with PBST for 15 minutes at RT 4 times. Afterwards, slides were incubated with secondary antibody solution (1:500 goat anti-mouse IgG Alexa Fluor 555 and 1X Hoechst diluted in blocking buffer) for 1 hour at RT. Slides were then washed with PBST for 15 minutes at RT 4 times and mounted in Vectashield mounting media.

For human, salamander, and killifish, the slides were thawed and dried at room temperature followed by immediate fixation with 4% PFA. The slides were washed in PBST (0.1% Tween-20 in PBS) 3 times for 5 min each wash at RT. Traces of PFA were quenched using 0.1M Glycine wash followed by 3 PBST washes separated by 5 minutes at RT. The sections were then permeabilized with 0.3% Triton X-100 for 15-20minutes and then blocked in blocking buffer (3% BSA, 5% Normal Goat Serum, 0.1% Triton X-100) for 1 hour at RT. The samples were then incubated with an anti-PAX7 (DSHB) primary antibody at 1:20 (for Killifish, Anti-PAX7, Santa Cruz, 1:100) and an anti-Laminin antibody at 1:1000 dilution overnight for 12-16 hours at 4°C. The following day, slides were washed for 15 minutes in PBST 4 times at RT, followed by incubation with secondary antibodies diluted in blocking buffer for 1 hour at RT. Hoechst was added for the detection of nuclei. Slides were washed in PBST 4 times, for 15 min each, and mounted using Vectashield mounting media and imaged.

##### Microscopy

Whole muscle sections of all species were imaged at 20X magnification using the Zeiss AxioScan.Z1 slide scanner. AxioScan yields a “scene” per sample, which is one large, stitched image of all the sections on the slide. Images of each individual section were obtained by splitting the scenes in Zeiss ZEN Lite software. Representative images were taken at 20X magnification for mouse, human and axolotl samples. For Killifish, we took images at 40X magnification (with oil immersion) and using Z-stacking with a Zeiss LSM 800 AxioObserver.M2 upright confocal microscope. Maximum projection of Z-stack images was processed in Fiji.

##### Fiji processing and analysis

Images of each muscle section were loaded into Fiji where brightness and contrast were adjusted until the intensities of PAX7 and Hoechst signals were equivalent. A composite image was then generated and saved as a jpeg or TIFF file in preparation for Ilastik analysis. The area of each muscle section was measured by tracing the muscle section with the Wand tool and measuring the area (μm^2^) inside the selection with the Measure plugin. Area measurements in μm^2^ were converted to mm^2^ in Excel. Representative images were loaded into Fiji where brightness and contrast were adjusted to appropriate levels and the composite images were generated.

##### Manual Quantifications

Manual quantifications were conducted by three trained counters from the laboratory, all of whom were blinded when quantifying. Manual counters were given composite images of whole muscle sections of every species, stained for PAX7 and Hoechst – these images were the same images analyzed by Ilastik. Manual counters opened images in Fiji, where they added a grid to the image (to make counting more efficient and organized) and counted cells using the Cell Counter plug-in. Raw cell counts were then inputted into an excel sheet, where PAX7+ cells/mm^2^ was calculated by dividing the total cell count by the area (mm^2^) of the respective muscle section. Quantifications were unblinded by a separate lab member.

##### Ilastik Analysis

Composite images of whole muscle sections that were pre-processed in Fiji were next inputted into Ilastik. One Ilastik project was created per species used in this study. Images for killifish were very large so we split them into 4 smaller images to help Ilastik run more efficiently. The Ilastik workflow is comprised of two processes: Pixel Classification and Object Classification. Briefly, we will describe specific parameters used in the present analyses. For Pixel Classification, all features, Color/Intensity, Edge and Texture from 0.3-10 pixels were selected for training. We trained Ilastik to count PAX7+ cells on 3 images per species (> 50-100 cells per training). For Object Classification, the following size filters and thresholding parameters were used: 1) Mouse no injury (Method: simple, Input: 0, Smooth: 1.0-1.0, Threshold: 0.8, Size Filter Min: 10, Size Filter Max: 1000000); Mouse 7 dpi (Method: simple, Input: 0, Smooth: 1.0-1.0, Threshold: 0.5, Size Filter Min: 10, Size Filter Max: 1000000); Human (Method: simple, Input: 0, Final: 0, Smooth: 1.0-1.0, Threshold: 0.8, Size Filter Min: 10, Size Filter Max: 1000000); Axolotl no injury (Method: hysteresis, Input: 0, Final:0, Smooth: 1.0-1.0, Threshold Core:0.7, Threshold final: 0.6, Size Filter Min: 7, Size Filter Max: 1000000); Axolotl 14dpa (Method: hysteresis, Input: 0, Final:0, Smooth: 1.0-1.0, Threshold Core:0.7, Threshold final: 0.6, Size Filter Min: 7, Size Filter Max: 1000000); turquoise killifish young (Method: hysteresis, Input: 0, Smooth: 1.0-1.0, Threshold Core:0.4, Threshold final: 0.3, Size Filter Min: 5, Size Filter Max: 300); turquoise killifish old (Method: hysteresis, Input: 0, Smooth: 1.0-1.0, Threshold Core:0.4, Threshold final: 0.3, Size Filter Min: 3, Size Filter Max: 300). Once each Ilastik project was finalized, non-training images were analyzed by batch processing. The raw cell counts (in the form of csv files) per image were placed into an excel sheet, where PAX7+ cells/mm^2^ was calculated by dividing the total cell count by the area (mm^2^) of the respective muscle section.

### Quantification and statistical analysis

Statistical analysis was performed using GraphPad Prism. The exact number of N and the statistical test used for each experiment are noted in the figure legend. P-values less than 0.05 were considered statistically significant (* denotes a p-value σ; 0.05 and ** denotes a p-value σ; 0.01). NS means not significant (p-value > 0.05) in our statistical tests

## References

1. Fuchs E, Blau HM. Tissue stem cells: Architects of their niches. Cell Stem Cell. 2020;27(4):532–556. doi: 10.1016/j.stem.2020.09.011.

2. Kim S, Roh J, Park C. Immunohistochemistry for pathologists: Protocols, pitfalls, and tips. J Pathol Transl Med. 2016;50(6):411–418. doi: 10.4132/jptm.2016.08.08.

3. Mebratie DY, Dagnaw GG. Review of immunohistochemistry techniques: Applications, current status, and future perspectives. Semin Diagn Pathol. 2024;41(3):154–160. doi: 10.1053/j.semdp.2024.05.001.

4. Schindelin J, Arganda-Carreras I, Frise E, et al. Fiji: An open-source platform for biological-image analysis. Nat Methods. 2012;9(7):676–682. doi: 10.1038/nmeth.2019.

5. Schindelin J, Rueden CT, Hiner MC, Eliceiri KW. The ImageJ ecosystem: An open platform for biomedical image analysis. Mol Reprod Dev. 2015;82(7-8):518–529. doi: 10.1002/mrd.22489.

6. Errington TM, Denis A, Perfito N, Iorns E, Nosek BA. Challenges for assessing replicability in preclinical cancer biology. Elife. 2021;10:10.7554/eLife.67995. doi: 10.7554/eLife.67995.

7. Diaba-Nuhoho P, Amponsah-Offeh M. Reproducibility and research integrity: The role of scientists and institutions. BMC Res Notes. 2021;14(1):451–3. doi: 10.1186/s13104-021-05875-3.

8. Baker M. 1,500 scientists lift the lid on reproducibility. Nature. 2016;533(7604):452–454. doi: 10.1038/533452a.

9. Baker M, Dolgin E. Cancer reproducibility project releases first results. Nature. 2017;541(7637):269–270. doi: 10.1038/541269a.

10. Stirling DR, Carpenter AE, Cimini BA. CellProfiler analyst 3.0: Accessible data exploration and machine learning for image analysis. Bioinformatics. 2021;37(21):3992–3994. doi: 10.1093/bioinformatics/btab634.

11. Korber N. MIA is an open-source standalone deep learning application for microscopic image analysis. Cell Rep Methods. 2023;3(7):100517. doi: 10.1016/j.crmeth.2023.100517.

12. Stringer C, Wang T, Michaelos M, Pachitariu M. Cellpose: A generalist algorithm for cellular segmentation. Nat Methods. 2021;18(1):100–106. doi: 10.1038/s41592-020-01018-x.

13. Pachitariu M, Stringer C. Cellpose 2.0: How to train your own model. Nat Methods. 2022;19(12):1634–1641. doi: 10.1038/s41592-022-01663-4.

14. Berg S, Kutra D, Kroeger T, et al. Ilastik: Interactive machine learning for (bio)image analysis. Nat Methods. 2019;16(12):1226–1232. doi: 10.1038/s41592-019-0582-9.

15. Seale P, Ishibashi J, Scime A, Rudnicki MA. Pax7 is necessary and sufficient for the myogenic specification of CD45+:Sca1+ stem cells from injured muscle. PLoS Biol. 2004;2(5):E130. doi: 10.1371/journal.pbio.0020130.

16. Seale P, Sabourin LA, Girgis-Gabardo A, Mansouri A, Gruss P, Rudnicki MA. Pax7 is required for the specification of myogenic satellite cells. Cell. 2000;102(6):777–786. doi: 10.1016/s0092-8674(00)00066-0.

17. Seale P, Ishibashi J, Holterman C, Rudnicki MA. Muscle satellite cell-specific genes identified by genetic profiling of MyoD-deficient myogenic cell. Dev Biol. 2004;275(2):287–300. doi: 10.1016/j.ydbio.2004.07.034.

18. von Maltzahn J, Jones AE, Parks RJ, Rudnicki MA. Pax7 is critical for the normal function of satellite cells in adult skeletal muscle. Proc Natl Acad Sci U S A. 2013;110(41):16474– 16479. doi: 10.1073/pnas.1307680110.

19. Lepper C, Partridge TA, Fan C. An absolute requirement for Pax7-positive satellite cells in acute injury-induced skeletal muscle regeneration. Development. 2011;138(17):3639–3646. doi: 10.1242/dev.067595.

20. Sambasivan R, Yao R, Kissenpfennig A, et al. Pax7-expressing satellite cells are indispensable for adult skeletal muscle regeneration. Development. 2011;138(17):3647–3656. doi: 10.1242/dev.067587.

21. Shea KL, Xiang W, LaPorta VS, et al. Sprouty1 regulates reversible quiescence of a self-renewing adult muscle stem cell pool during regeneration. Cell Stem Cell. 2010;6(2):117–129. doi: 10.1016/j.stem.2009.12.015.

22. Mademtzoglou D, Geara P, Mourikis P, Relaix F. Pax7 haploinsufficiency impairs muscle stem cell function in cre-recombinase mice and underscores the importance of appropriate controls. Stem Cell Res Ther. 2023;14(1):294–1. doi: 10.1186/s13287-023-03506-1.

23. Arenas Gomez CM, Echeverri K. Salamanders: The molecular basis of tissue regeneration and its relevance to human disease. Curr Top Dev Biol. 2021;145:235–275. doi: 10.1016/bs.ctdb.2020.11.009.

24. Riquelme-Guzman C, Sandoval-Guzman T. The salamander limb: A perfect model to understand imperfect integration during skeletal regeneration. Biol Open. 2024;13(2):bio060152. doi: 10.1242/bio.060152. Epub 2024 Feb 6. doi: 10.1242/bio.060152.

25. Boluk A, Yavuz M, Demircan T. Axolotl: A resourceful vertebrate model for regeneration and beyond. Dev Dyn. 2022;251(12):1914–1933. doi: 10.1002/dvdy.520.

26. McCusker C, Gardiner DM. The axolotl model for regeneration and aging research: A mini-review. Gerontology. 2011;57(6):565–571. doi: 10.1159/000323761.

27. Boluk A, Yavuz M, Demircan T. Axolotl: A resourceful vertebrate model for regeneration and beyond. Dev Dyn. 2022;251(12):1914–1933. doi: 10.1002/dvdy.520.

28. McCusker C, Gardiner DM. The axolotl model for regeneration and aging research: A mini-review. Gerontology. 2011;57(6):565–571. doi: 10.1159/000323761.

29. Tajer B, Savage AM, Whited JL. The salamander blastema within the broader context of metazoan regeneration. Front Cell Dev Biol. 2023;11:1206157. doi: 10.3389/fcell.2023.1206157.

30. Fei J, Schuez M, Knapp D, Taniguchi Y, Drechsel DN, Tanaka EM. Efficient gene knockin in axolotl and its use to test the role of satellite cells in limb regeneration. Proc Natl Acad Sci U S A. 2017;114(47):12501–12506. doi: 10.1073/pnas.1706855114.

31. Morrison JI, Loof S, He P, Simon A. Salamander limb regeneration involves the activation of a multipotent skeletal muscle satellite cell population. J Cell Biol. 2006;172(3):433–440. doi: 10.1083/jcb.200509011.

32. Hu C, Brunet A. The african turquoise killifish: A research organism to study vertebrate aging and diapause. Aging Cell. 2018;17(3):e12757. doi: 10.1111/acel.12757.

33. de Bakker DEM, Valenzano DR. Turquoise killifish: A natural model of age-dependent brain degeneration. Ageing Res Rev. 2023;90:102019. doi: 10.1016/j.arr.2023.102019.

34. Kim Y, Nam HG, Valenzano DR. The short-lived african turquoise killifish: An emerging experimental model for ageing. Dis Model Mech. 2016;9(2):115–129. doi: 10.1242/dmm.023226.

35. Kim Y, Nam HG, Valenzano DR. The short-lived african turquoise killifish: An emerging experimental model for ageing. Dis Model Mech. 2016;9(2):115–129. doi: 10.1242/dmm.023226.

36. Graf M, Hartmann N, Reichwald K, Englert C. Absence of replicative senescence in cultured cells from the short-lived killifish nothobranchius furzeri. Exp Gerontol. 2013;48(1):17–28. doi: 10.1016/j.exger.2012.02.012.

37. Ruparelia AA, Salavaty A, Barlow CK, et al. The african killifish: A short-lived vertebrate model to study the biology of sarcopenia and longevity. Aging Cell. 2024;23(1):e13862. doi: 10.1111/acel.13862.

38. Teefy BB, Adler A, Xu A, Hsu K, Singh PP, Benayoun BA. Dynamic regulation of gonadal transposon control across the lifespan of the naturally short-lived african turquoise killifish. Genome Res. 2023;33(1):141–153. doi: 10.1101/gr.277301.122.

39. Mayeuf-Louchart A, Hardy D, Thorel Q, et al. MuscleJ: A high-content analysis method to study skeletal muscle with a new fiji tool. Skelet Muscle. 2018;8(1):25–0. doi: 10.1186/s13395-018-0171-0.

40. Danckaert A, Trignol A, Le Loher G, et al. MuscleJ2: A rebuilding of MuscleJ with new features for high-content analysis of skeletal muscle immunofluorescence slides. Skelet Muscle. 2023;13(1):14–1. doi: 10.1186/s13395-023-00323-1.

41. Smith LR, Barton ER. 4417508; SMASH - semi-automatic muscle analysis using segmentation of histology: A MATLAB application. Skelet Muscle. 2014;4:21. http://www.ncbi.nlm.nih.gov/pubmed/25937889. doi: 10.1186/2044-5040-4-21.

42. Reyes-Fernandez PC, Periou B, Decrouy X, Relaix F, Authier FJ. Automated image-analysis method for the quantification of fiber morphometry and fiber type population in human skeletal muscle. Skelet Muscle. 2019;9(1):15–7. doi: 10.1186/s13395-019-0200-7.

43. Kastenschmidt JM, Ellefsen KL, Mannaa AH, et al. QuantiMus: A machine learning-based approach for high precision analysis of skeletal muscle morphology. Front Physiol. 2019;10:1416. doi: 10.3389/fphys.2019.01416.

44. Almada AE, Wagers AJ. PMC4918817; molecular circuitry of stem cell fate in skeletal muscle regeneration, ageing and disease. Nat Rev Mol Cell Biol. 2016;17(5):267–279. https://www.ncbi.nlm.nih.gov/pubmed/26956195. doi: 10.1038/nrm.2016.7.

