## Supplementary Figure 1 for "Ilastik: a machine learning image analysis platform to interrogate stem cell fate decisions across multiple vertebrate species"

A

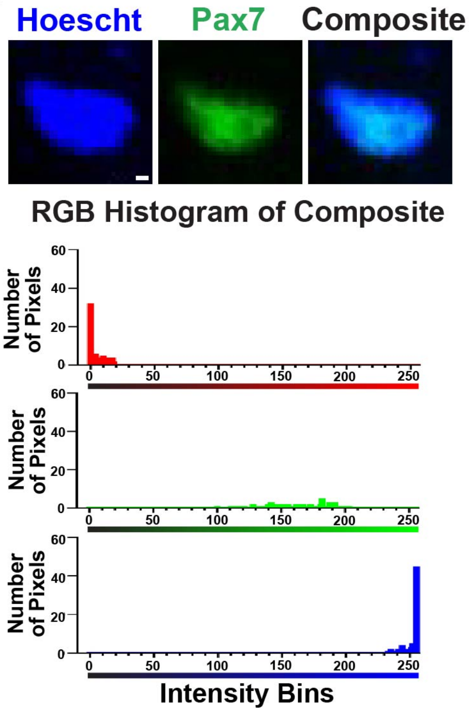


**Supplemental Figure 1.**
(A) Example of one cell displaying an inadequate adjustment to brightness/contrast (B/C) of nuclear stain (Hoescht in blue) and PAX7 (Green) (top). Representative histogram showing what an inadequate B/C adjustment looks like when looking at pixel intensity of the red, green, and blue channels (bottom). Scale bar represents 1um.
